# Serological fingerprints link antiviral activity of therapeutic antibodies to affinity and concentration

**DOI:** 10.1101/2022.02.03.478946

**Authors:** Sebastian Fiedler, Sean R. A. Devenish, Alexey S. Morgunov, Alison Ilsley, Francesco Ricci, Marc Emmenegger, Vasilis Kosmoliaptsis, Elitza S. Theel, John R. Mills, Anton M. Sholukh, Adriano Aguzzi, Akiko Iwasaki, Andrew K. Lynn, Tuomas P. J. Knowles

**Affiliations:** Fluidic Analytics, Unit A, The Paddocks Business Centre, Cherry Hinton Road, Cambridge CB1 8DH, United Kingdom; Centre for Misfolding Diseases, Yusuf Hamied Department of Chemistry, University of Cambridge, Lensfield Road, Cambridge CB2 1EW, United Kingdom; Institute of Neuropathology, University of Zurich, 8091 Zurich, Switzerland; Department of Surgery, University of Cambridge, Addenbrookes Hospital, Cambridge, CB2 0QQ, United Kingdom; NIHR Blood and Transplant Research Unit in Organ Donation and Transplantation, University of Cambridge, Hills Road, Cambridge CB2 0QQ, United Kingdom; NIHR Cambridge Biomedical Research Centre, Hills Road, Cambridge CB2 0QQ, United Kingdom; Division of Clinical Microbiology, Department of Laboratory Medicine and Pathology, Mayo Clinic, Rochester, Minnesota, USA; Department of Laboratory Medicine and Pathology, Mayo Clinic, Rochester, Minnesota, USA; Center for MS and Autoimmune Neurology, Mayo Clinic, Rochester, Minnesota, USA; Vaccine and Infectious Disease Division, Fred Hutchinson Cancer Research Center, Seattle, Washington, USA; Department of Immunobiology, Yale School of Medicine, New Haven, CT 06519, USA; Department of Epidemiology of Microbial Diseases, Yale School of Public Health, New Haven, CT 06510, USA; Department of Molecular, Cellular and Developmental Biology, Yale University, New Haven, CT 06511, USA; Howard Hughes Medical Institute, Chevy Chase, MD 20815, USA; Cavendish Laboratory, Department of Physics, University of Cambridge, JJ Thomson Ave, Cambridge CB3 0HE, United Kingdom

**Keywords:** Omicron, variants of concern, monoclonal antibodies, humoral immunity, antibody affinity, microfluidics

## Abstract

We assessed the affinities of the therapeutic monoclonal antibodies (mAbs) cilgavimab, tixagevimab, sotrovimab, casirivimab, and imdevimab to the receptor binding domain (RBD) of wild type, Delta, and Omicron spike. The Omicron RBD affinities of cilgavimab, tixagevimab, casirivimab, and imdevimab decreased by at least two orders of magnitude relative to their wild type equivalents, whereas sotrovimab binding was less severely impacted. These affinity reductions correlate with reduced antiviral activities of these antibodies, suggesting that simple affinity measurements can serve as an indicator for activity before challenging and time-consuming virus neutralization assays are performed. We also compared the properties of these antibodies to serological fingerprints (affinities and concentrations) of wild type RBD specific antibodies in 74 convalescent sera. The affinities of the therapeutic mAbs to wild type and Delta RBD were in the same range as the polyclonal response in the convalescent sera indicative of their high antiviral activities against these variants. However, for Omicron RBD, only sotrovimab retained affinities that were within the range of the polyclonal response, in agreement with its high activity against Omicron. Serological fingerprints thus provide important context to affinities and antiviral activity of mAb drugs and could guide the development of new therapeutics.

## Introduction

The SARS-CoV-2 B.1.1.529 variant (Omicron) was first reported in Botswana and in South Africa in November 2021 and was classified as a variant of concern by the world health organization (WHO) on 26^th^ of November 2021^1,2^. By mid-December 2021, the Omicron variant was detected in more than 30 countries and by late January 2022 was the dominant lineage worldwide.

The Omicron variant is characterized by a large number of mutations present in the spike and nucleocapsid proteins. Most critical for viral fitness and immune evasion are likely 34 mutations within the Omicron spike protein with 10 mutations within the N-terminal domain, 15 in the receptor binding domain (RBD), 3 related to the furin cleavage site and 6 in the S2 region (Tables S1 and S2). Of these mutations, 13 had been observed in previous variants of SARS-CoV-2 but never in a single lineage, as summarized in Tables S1 and S2. Despite this large number of mutations, Omicron still utilizes angiotensin converting enzyme (ACE2) as host receptor and binds with similar affinity than the original Wuhan strain (referred to as wild type throughout the paper)^3,4^.

Omicron mutations reduce the virus neutralization efficacy of some approved or clinical-stage antibody drugs. Casirivimab/imdevimab (Regeneron) and bamlanivimab/etesevimab (Lilly) lose their ability to neutralize, while cilgavimab/tixagevimab (AstraZeneca) and sotrovimab (GSK) retain some degree of efficacy^3,5–9^.

Virus neutralization of Omicron is also strongly reduced in convalescent sera from patients infected with prior lineages and in the sera of double-vaccinated individuals who had been vaccinated with BNT162b2, mRNA-1273, Ad26.COV2.S, ADZ1222, Sputnik V, or BBIBP-CorV^3,6,9–11^. Triple vaccinated individuals who have received BNT162b2 or mRNA-1273 also show reduced neutralization efficacy against Omicron relative to wild type and Delta, although the retained efficacies are considerably higher than for convalescent or double-vaccinated individuals^3,6,8,9^. Also, individuals who have been infected with either Delta or an earlier variant of SARS-CoV-2 and subsequently been vaccinated retained considerable titers of neutralizing antibodies (NAbs)^3,8,9^.

The ability of the Omicron variant to evade humoral immune responses, whether induced by infection or vaccination, is expected to cause more reinfections and breakthrough infections. Despite reports of a higher proportion of Omicron infections leading to milder disease outcomes^12–17^, very high case numbers resulting from a more transmissible Omicron variant would still pose a significant public health risk.

Here, we used microfluidic diffusional sizing (MDS)^18–21^ to measure the in-solution binding affinities to the spike RBD of the wild-type, Delta and Omicron variants of five therapeutic monoclonal antibodies (casirivimab, imdevimab, sotrovimab, tixagevimab and cilgavimab) administered to reduce SARS-CoV-2 viral load and alleviate COVID-19 symptoms. All five antibodies bind wild type and Delta SARS-CoV-2 spike with high affinities and are potent virus neutralizing agents^22^. For four of these five drugs, the affinity for the Omicron spike RBD was more than two orders of magnitude lower than the affinity for the wild type spike RBD; by contrast, sotrovimab retained significantly higher affinity for the Omicron spike RBD. The MDS-based antibody affinities determined were consistent with published virus-neutralization IC_50_ values and provide a quantitative explanation for the relative efficacies of all five antibodies against Omicron. We also considered the affinities of these five mAbs to spike RBDs of the wild type, Delta, and Omicron variants in the context of antibody fingerprints, consisting of affinity and concentration data, obtained from the sera of COVID-19 convalescents by microfluidic antibody affinity profiling (MAAP)^18–21^. MAAP showed that wild type and Delta spike RBD affinities of all five therapeutic antibodies were within the affinity range typical of polyclonal anti-wild type spike RBD antibodies generated by the humoral immune response. Against Omicron spike RBD, only sotrovimab stayed within this affinity range, while the other four mAbs had affinities several orders of magnitude lower than antibodies found in convalescent sera. We also utilized MAAP to fingerprint the antibody response against wild type, Delta and Omicron spike RBDs in the working reagent for anti-SARS-CoV-2 immunoglobulin^23^, a pooled plasma standard from individuals who recovered from COVID-19 in 2020. We observed considerable cross-reactivity to Omicron spike RBD and roughly half the concentration of binding antibodies as compared with wild type and Delta.

## Results and Discussion

The binding affinities of cilgavimab, tixagevimab, sotrovimab, casirivimab and imdevimab to Omicron spike RBD, Delta spike RBD, and wild type spike RBD were determined by MDS (Figure 1). To do so, equilibrium binding curves were acquired by titrating each antibody against constant concentrations of each spike RBD. The formation of RBD–antibody complexes was monitored based on an increase of the hydrodynamic radius (*R*_h_) of the RBD species in solution.

**Figure 1.**
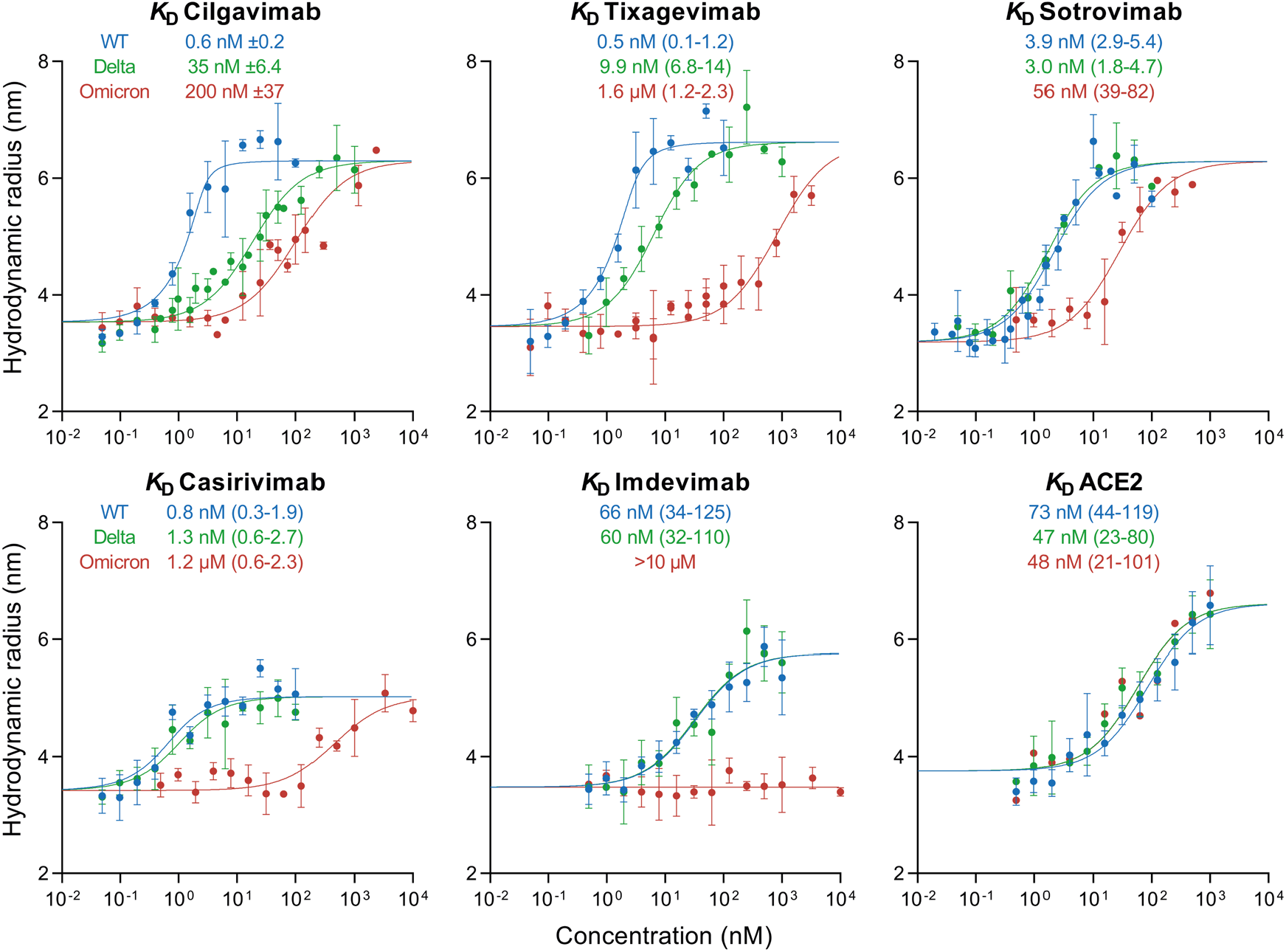
Equilibrium binding curves of cilgavimab, tixagevimab, sotrovimab, casirivimab, imdevimab, and ACE2 binding to spike RBD proteins from SARS-CoV-2 wild type (blue) as well as variants Delta (green) and Omicron (red) as determined by microfluidic diffusional sizing (MDS). Error bars are standard deviations from triplicate measurements. *K*_D_ values are best fits with standard errors from non-linear least squares fits in terms of a 2:1 binding model (antibodies) or a 1:1 binding model (ACE2).

At the lowest antibody concentrations, in the absence of binding, wild type, Delta, and Omicron spike RBDs displayed *R*_h_ values of 3.40 nm (SD = 0.27 nm), as observed previously^18,20,21^.

Notably, the *K*_D_ values measured here using microfluidic diffusional sizing are in agreement with published values obtained by SPR^3,4,22^.

Therapeutic antibody binding to Omicron RBD was previously reported in a study using BLI, which detected no binding for imdevimab, consistent with the MDS results found here. By contrast, however, the BLI results in this same study reported a low nanomolar affinity to Omicron for cilgavimab and sub-nanomolar affinities for tixagevimab, sotrovimab, and casirivimab^5^ – results that differ strongly from the affinity values determined by MDS in our present study that show reduced affinity of all four of these mAbs for Omicron. Given the density of mutations in Omicron it seems likely that the affinity determined by BLI is artefactual, a possibility supported by the observation of lost neutralization efficacy for most of these antibodies when challenged with the Omicron variant^3,5–9^.

As shown in Figure 2, the five antibodies tested have different epitopes on the RBD depending on their modes of action. Cilgavimab and tixagevimab are used as a combination drug, bind to non-overlapping epitopes, and inhibit ACE2 binding^27–29^. Cilgavimab binds both the up and down conformation of RBD while tixagevimab exclusively binds to the up confirmation. For cilgavimab, the substitutions N440K and G446S are likely to affect binding and neutralization. Tixagevimab binds to the left shoulder of RBD with Omicron RBD mutations S477N, T478K, and E484A likely to interfere with binding and neutralization. Sotrovimab binds outside the receptor binding motif to a site that involves the N343-linked glycan^30,31^. This epitope might be sensitive to G339D and N440K substitutions in the Omicron spike. Casirivimab and imdevimab prevent viral spike proteins from binding to ACE2^22^. The epitope of casirivimab overlaps with the ACE2 binding site, whereas imdevimab binds on the side of the RBD and sterically blocks ACE2 from accessing the spike protein. The binding epitopes of both antibodies contain Omicron mutations: for casirivimab the mutations K417N, E484A, Q493R are all within 5 Å of the antibody, while imdevimab is within 5 Å of the mutations N440K and Q498R (Figure 2).

**Figure 2.**
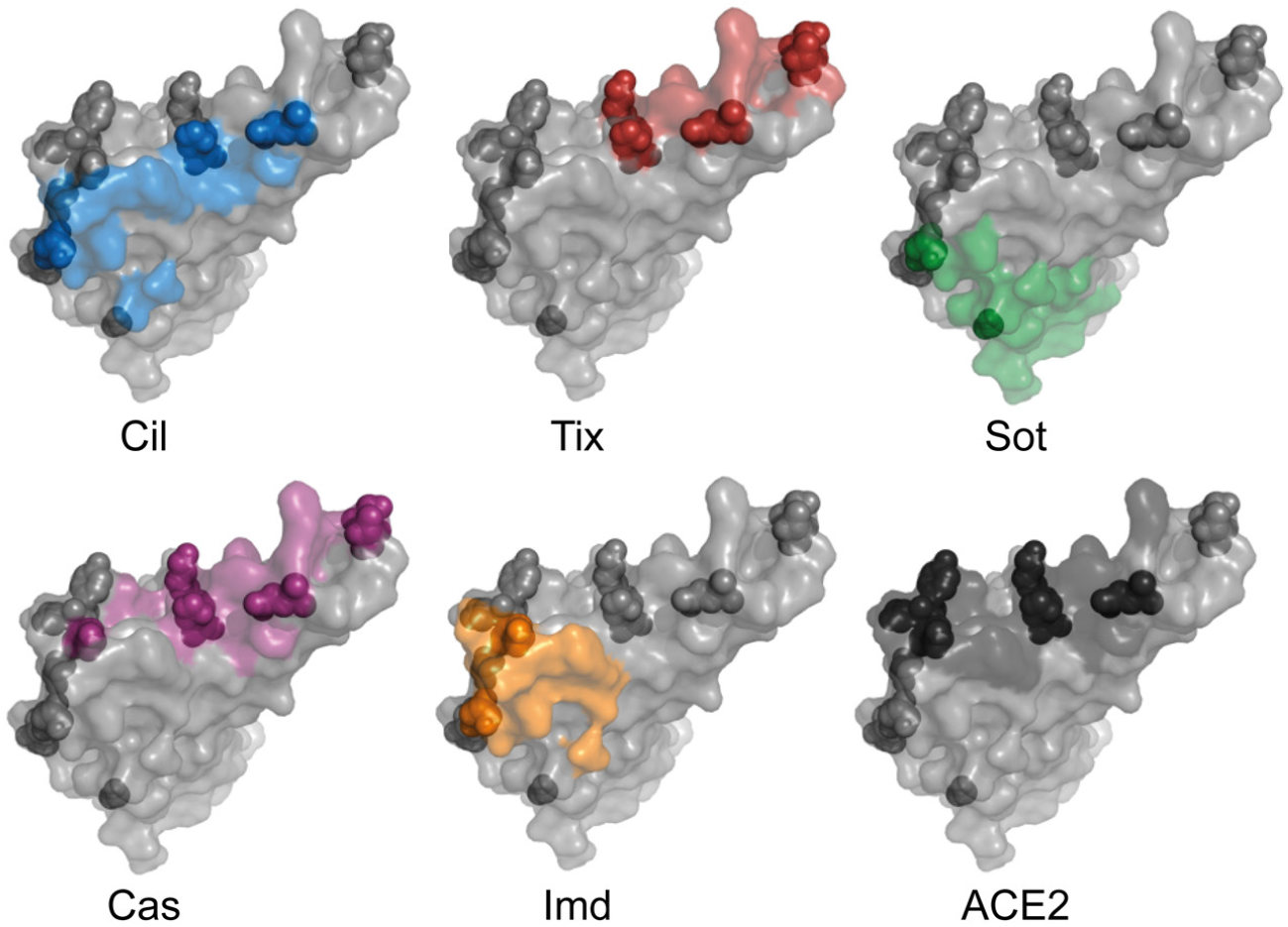
Omicron spike mutations compared to antibody epitopes and the ACE2 binding motif. RBD is shown as grey surface with residues within 5 Å of indicated interacting partner colored and Omicron mutations shown as spheres. PDB codes for structures are: 7L7E^24^ (cilgavimab and tixagevimab), 7SOC^25^ (sotrovimab), 6XDG^22^ (casirivimab and imdevimab), and 7DQA^26^ (ACE2).

Relative to wild type RBD, only cilgavimab and tixagevimab showed a reduced binding affinity to Delta RBD (Figure 3A). For Omicron RBD, cilgavimab, tixagevimab, casirivimab, and imdevimab displayed binding affinities reduced by at least two orders of magnitude. Due to its highly conserved epitope^30^, the affinity of sotrovimab to Omicron RBD was reduced by only a factor of 10.

**Figure 3.**
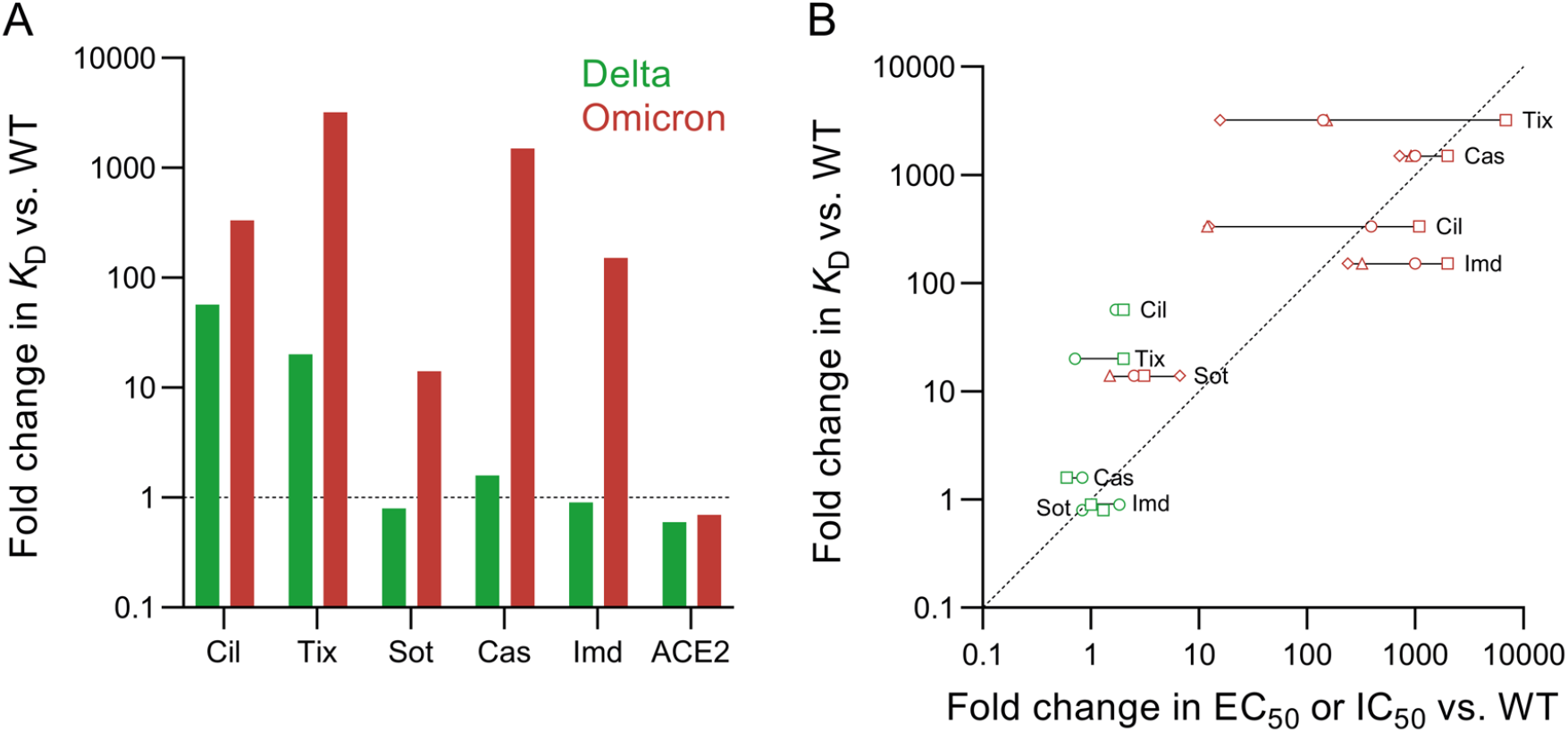
(A) Affinity (*K*_D_) changes of therapeutic COVID-19 antibodies and ACE2 in the presence of Delta (green) and Omicron (red) spike RBDs as compared with wild type RBD. (B) *K*_D_ changes of therapeutic COVID-19 antibodies correlated with their published half-maximal effective concentrations (EC_50_) or half-maximal inhibitory concentrations (IC_50_) of focus reduction neutralization tests (FRNT; live virus) or pseudovirus neutralization tests (PNT), respectively. Triangles and diamonds are FRNTs using VERO-TMPRSS2 and VERO-hACE2-TMPRSS2 cells, respectively, taken from VanBlargan et al.^7^. Squares and circles are PNTs taken from Cao et al. ^5^ and Liu et al.^6^, respectively.

The affinities of both Delta and Omicron RBD for ACE2 are very similar to that of wild type RBD (Figure 3A). In the case of Delta RBD, this would be expected as this variant does not carry any changes in the ACE2 binding interface^4^. Omicron RBD, on the other hand, has eight substitutions in the ACE2 binding interface^4^, so it is surprising that the binding affinity is not impacted. These data highlight the critical role this interaction plays in the viral life cycle and hence the selective pressure on RBD mutants to maintain efficient ACE2 binding. Given the retained ACE2 affinity of the Omicron variant, cilgavimab, tixagevimab, and casirivimab, which inhibit viral ACE2 binding, would require extremely high concentrations to achieve relevant levels of inhibition due to their strongly reduced affinity.

The changes in affinities, relative to the wild type spike RBD, of these five antibodies that result from the Delta and Omicron mutations suggest close correlations between in-solution binding affinity measurements and changes in antiviral activity. As shown in Figure 3B, our in-solution affinity measurements reflect changes in EC_50_ and IC_50_ obtained by focus reduction neutralization tests and pseudovirus neutralization assays^3,5–9^ very well. For example, each of casirivimab, imdevimab, and sotrovimab have similar changes in affinities and in EC_50_/IC_50_ values when challenged with wild type or Delta RBDs. When challenged with the Omicron variant, all tested antibodies except sotrovimab experience strong reduction in affinity in line with strong decrease in IC_50_ values. Sotrovimab retains both considerable binding affinity and neutralization efficacy. The correlation of in-solution *K*_D_ and EC_50_/IC_50_ values raises the prospect of a straight-forward method for predicting antiviral activity. MDS provides universally comparable results in the form of absolute affinities (*K*_D_), requires less than 1 hour, uses less than 4 µg of antibody, and is simple to perform as no cell culture or handling of live viruses is required. In addition to being much more time consuming and complex experiments, virus neutralization assays can yield variable results depending, for example, on the exact type of assay that is used (Figure 3B). Here, MDS can serve as a complementary method to support observations made in more complex biological systems.

Next, we analyzed how the spike RBD binding affinities of cilgavimab, tixagevimab, sotrovimab, casirivimab, and imdevimab compared to polyclonal antibody responses in unvaccinated COVID-19 convalescent individuals (Figure 4). We were interested to assess the behavior of these monoclonal antibody therapies in the context of the humoral response that results from infection. Generally, affinities of serum antibodies and their specific concentrations are difficult to measure. To address this issue, we recently introduced microfluidic affinity profiling (MAAP)^18,20,21,32^ as an advanced serological assay which utilizes MDS to provide a fingerprint of the antibodies able to bind an antigen probe. Specifically, the fingerprint consists of the in-solution *K*_D_ and the concentration of antibody binding sites in any complex biological background. To assess the functional immune response against SARS-CoV-2, MAAP was used to determine the antibody affinity against fluorescently labeled wild type RBD as well as the concentration of the binding antibodies directly in biological samples. This granular, quantitative view on the functional immune response of individuals is advantageous over commonly measured antibody titers which are a combination of both affinity and concentration. While a high titer can be achieved through either a low concentration of high affinity antibodies or a high concentration of low affinity antibodies, MAAP can readily distinguish these physiologically very distinct scenarios.

**Figure 4.**
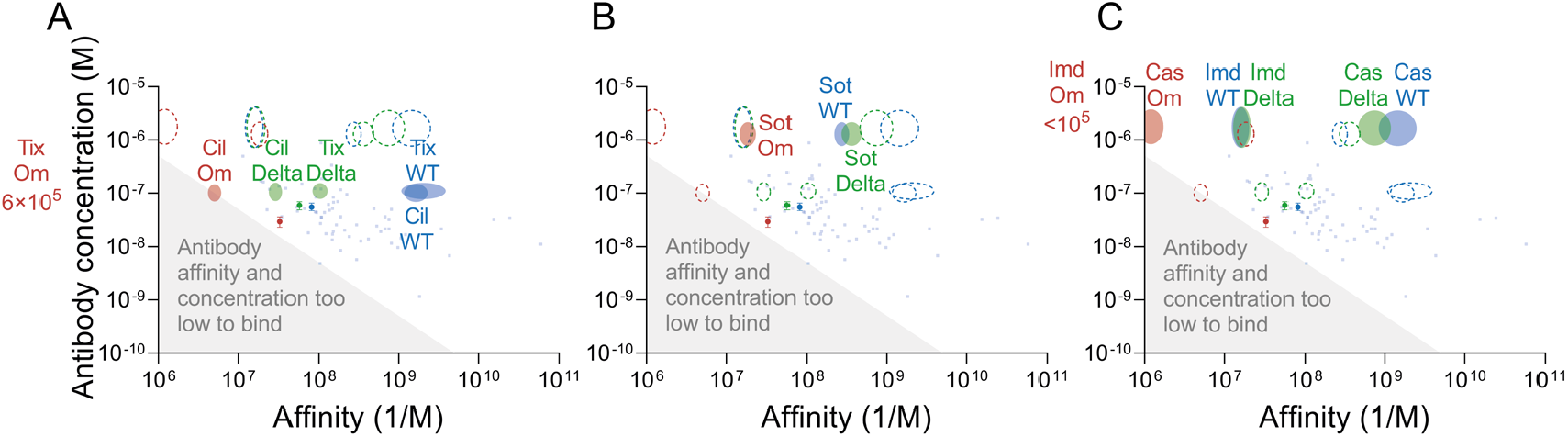
Affinities and average maximum post-dosage concentrations of therapeutic antibodies (A: cilgavimab and tixagevimab, B: sotrovimab, and C: casirivimab and imdevimab) in comparison with affinities and concentrations of COVID-19 convalescent serum or plasma. The width and height of the monoclonal antibody ovals are the standard error affinities (*K*_A_ determined by MDS) against wild type spike RBD (blue), Delta spike RBD (green), or Omicron spike RBD (red) and maximum serum concentrations obtained after dosage^33–35^ (average ± 1 SD), respectively. Light blue squares correspond to microfluidic antibody affinity profiling (MAAP) fingerprints of convalescent serum samples collected from various cohorts of unvaccinated individuals using wild type spike RBD as antigen (see Table S3 for details). Circles correspond to MAAP performed on the working reagent for anti-SARS-CoV-2 immunoglobulin^23^ (pooled COVID-19 convalescent plasma) using wild type spike RBD (blue), Delta spike RBD (green), or Omicron spike RBD (red) as antigens. The shaded gray region of the plot indicates the area where antibody concentration is less than *K*_D_/2 such that binding is not able to exceed 50%.

Figure 4 collates 74 MAAP results performed by us using samples of COVID-19 convalescent serum or plasma from unvaccinated individuals and wild type RBD as the antigen (see Table S3 for details). As shown previously^20^, both affinity and concentration of antibodies vary over several orders of magnitude between individuals. However, 75% of samples contained antibodies with affinities greater than 10^8^ nM^-1^ (*K*_D_ ≤10 nM) and total binding site concentrations ≤150 nM (Figure 4). Cilgavimab, tixagevimab, casirivimab and imdevimab are derived from convalescent individuals that had been exposed to wild type RBD early in the pandemic^22^. Sotrovimab was obtained from a patient who was exposed to the SARS virus during the early 2000s^36^. As might be expected, the affinities for wild type RBD of the monoclonal antibodies that compete directly with ACE2 binding (Figure 4A: cilgavimab, tixagevimab, and Figure 4C: casirivimab) are tighter than the majority of the population’s polyclonal antibody response to the same antigen. Sotrovimab (Figure 4B) and imdevimab (Figure 4C) are located in the same region of the plots indicative of slightly lower affinities than the typical polyclonal response.

The reduction in affinity of cilgavimab and tixagevimab against Delta brings them within the range of the polyclonal anti-wild type RBD antibody response produced by most convalescent individuals (Figure 4A). On the other hand, casirivimab, imdevimab and sotrovimab do not show a considerable shift when challenged with Delta RBD as their affinity is largely unaffected. Against Omicron, all antibodies, except sotrovimab, are either within or close to the regime for which less than 50% of target can be bound (grey area) and so would be expected to provide minimal therapeutic benefit.

Polyclonal anti-wild type RBD antibodies constitute a mixture of non-NAbs and NAbs, with NAbs varying in their neutralization mechanisms^5,37–39^. Regardless of their mechanisms, loss of NAb binding affinity correlates with viral immune escape^37–39^ (Figure 3B), as RBD binding is a pre-requisite for neutralization. In addition to the NAb affinity, the degree to which NAbs bind is determined by the viral load (i.e., the antigen concentration) and the NAb concentration. With reduced affinity, higher NAb concentrations are required to achieve the same levels of binding for a given viral load. If the affinity is too low, even very high NAb concentrations are not sufficient to achieve considerable binding. Since 60% of anti-RBD antibodies isolated from convalescent individuals showed neutralization^39^, a reduction in the average affinity of a polyclonal NAb mixture is likely linked to reduced virus neutralization.

We also assessed how the antibody affinity and the binding site concentration in a pooled convalescent standard plasma responds to Omicron spike RBD. For this experiment, we performed MAAP on the working reagent for anti-SARS-CoV-2 immunoglobulin, which is pooled plasma from COVID-19 convalescent individuals collected between April and May 2020^23^. Antibody affinities and concentrations against wild type RBD were well within the range observed for most of the individual samples (Figure 4). When challenged with Delta spike RBD, the affinity of antibodies in the pooled plasma decreased slightly by a factor of approximately 1.5 while the concentration of binding sites remained largely unchanged. Surprisingly, against Omicron spike RBD the antibody affinity and concentration were reduced by factors of just 2.5 and 2.0, respectively, compared to wild type. The binding site concentration is still 2-fold higher than *K*_D_ suggesting that these polyclonal antibody mixtures retain binding capabilities against Omicron spike RBD. This observation is in line with reports that a considerable population retain some degree of Omicron neutralization after infection with wild type or Alpha strains^9^. During infection, highly neutralizing antibodies with high affinity arise very quickly, such that just 2 mutations from germline can boost affinity 100-fold. However, the vast majority of induced antibodies against SARS-CoV-2 remain near-germline and thus can be more promiscuous for epitope mutations^37–39^. Figure 4 shows that compared with the therapeutic monoclonal antibodies, most individual sera and plasmas have rather low affinity and only very few demonstrate picomolar affinities. For such moderate affinity antibodies, a drop in affinity due to epitope changes is likely to be less profound than for antibodies with high affinity. For the latter, epitope mutation can be detrimental as evidenced by the profoundly reduced affinities of casirivimab, tixagevimab and cilgavimab, for example.

## Conclusions

Using microfluidic diffusional sizing, we quantified the binding affinities of five therapeutic antibodies to spike receptor binding domains of the wild type variant, the Delta variant, and the Omicron variant of SARS-CoV-2. The affinities of cilgavimab, tixagevimab, casirivimab, and imdevimab to Omicron spike RBD were reduced by several orders of magnitude, whereas sotrovimab retained considerable binding affinity. These affinity reductions in the presence of Omicron RBD agree very well with the reduced antiviral activity of these antibodies for which sotrovimab is also the least affected. These results suggest that simple in-solution affinity measurements can serve to evaluate antiviral activity before complex and time-consuming virus neutralization assays are performed. Serological fingerprints generated by microfluidic antibody affinity profiling of samples from COVID-19 convalescent individuals reveal serum-antibody affinity and concentration and provide further context to this finding. Out of the five tested antibodies, only sotrovimab retained affinities similar to those of polyclonal antibodies specific for wild type RBD, which is again indicative of its high antiviral activity against the Omicron variant. Our results represent a new way of linking monoclonal antibody affinities with their antiviral activity and serological fingerprints which has the potential to guide the development of new therapeutics to fit the affinity window of antibodies generated by the humoral immune response.

## Materials and Methods

### Spike RBD reagents

| Protein name | SARS-CoV-2 variant | Pango lineage | Vendor | Article number | Amino-acid sequence | Tags | Mutated amino-acid residues |
| --- | --- | --- | --- | --- | --- | --- | --- |
| Wild type spike RBD | Wuhan-Hu-1 sequence (NC_045512) | A | Sino Biological | 40592-V08H | R319-F541 | C-His |  |
| Delta spike RBD | Delta | B.1.617.2 | Sino Biological | 40592-V08H90 | R319-F541 | C-His | L452R, T478K |
| Omicron spike RBD | Omicron | B.1.1.529 | Sino Biological | 40592-V08H121 | R319-F541 | C-His | G339D, S371L, S373P, S375F, K417N, N440K, G446S, S477N, T478K, E484A, Q493R, G496S, Q498R, N501Y, Y505H |

### Antibody reagents

| Antibody name | Alternative name | Vendor | Article number |
| --- | --- | --- | --- |
| Tixagevimab Biosimilar | AZD8895 | ProteoGenix | PX-TA1032 |
| Cilgavimab Biosimilar | AZD1061 | ProteoGenix | PX-TA1033 |
| Sotrovimab Biosimilar | VIR-7831 | ProteoGenix | PX-TA1637 |
| Casirivimab | REGN10933 | ProteoGenix | PTXCOV-A552 |
| Imdevimab | REGN10987 | ProteoGenix | PTXCOV-A553 |

### Sample collection, ethics, and biosafety

All plasma and serum samples were collected from unvaccinated convalescent individuals. Unless otherwise stated, no information regarding the application of therapeutic antibodies, immunosuppressant drugs or other therapeutic agents was available for the purposes of this study. All such sample collection was performed in accordance with one of the following:

#### For all samples^20,40,41^ collected by University Hospital Zurich

All experiments and analyses involving samples from human donors were conducted with the approval of the local ethics committee (KEK-ZH-Nr.2015-0561, BASEC-Nr. 2018-01042, and BASEC-Nr. 2020-01731), in accordance with the provisions of the Declaration of Helsinki and the Good Clinical Practice guidelines of the International Conference on Harmonisation.

EDTA plasma from healthy donors and from convalescent individuals was obtained from the Blutspendedienst (BDS) Kanton Zürich from donors who signed the consent that their samples can be used for conducting research. Samples from patients with COVID-19 were collected at the University Hospital Zurich from patients who signed an informed consent.

#### For all samples collected by the Fred Hutchinson Cancer Research Center and the University of Washington

Informed consent was obtained from all human subjects. Plasma samples from SARS-CoV-2 seropositive individuals were obtained from a Fred Hutchinson Cancer Research Center repository and from the repository of the University of Washington. FHCRC repository was assembled from a COVID-19 seroepidemiology study conducted in a single county in the western US and the study was approved by the FHCRC institutional review board (#10453)^42^. UW repository was formed from Seattle-area participants recruited for potential donation of single-donor plasma units (ClinicalTrials.gov: NCT04338360), and plasma for manufacture of a pooled anti–SARS-CoV-2 product (NCT04344977)^43^.

#### For all samples purchased from BioIVT, LLC

BioIVT sought informed consent from each subject, or the subjects legally authorized representative, and appropriately documented this in writing. All samples are collected under IRB-approved protocols.

#### For all samples collected by Mayo Clinic

Samples were residual waste specimens that were fully deidentified and handled according to the policies for the use of waste specimens approved by the Mayo Clinic Institutional Review Board (IRB). The IRB review processes are based on the Federal Policy for the Protection of Human Subjects (“Common Rule”), the Belmont Report and provisions of 45CFR46 — “Protection of Human Subjects”.

### Fluorescent labeling of proteins

Recombinant proteins were labeled with Alexa Fluor(tm) 647 NHS ester (Thermo Fisher) as described previously^18^. In brief, solution containing 150 µg of spike RBD was mixed with dye at a three-fold molar excess in the presence of NaHCO_3_ (Merck) buffer at pH 8.3 and incubated at 4 °C overnight. Unbound label was removed by size-exclusion chromatography (SEC) on an ÄKTA pure system (Cytiva) using a Superdex 75 Increase 10/300 column (Cytiva). Labeled and purified proteins were stored at -80 °C in PBS pH 7.4 containing 10% (w/v) glycerol as cryoprotectant.

### Equilibrium binding measurements by microfluidic diffusional sizing (MDS)

Binding affinity of antibodies and spike RBD proteins was measured on a Fluidity One-M (Fluidic Analytics). Fluorescently labeled RBD at a concentration of 1 nM or 5 nM was mixed with unlabelled antibody at 12 different, decreasing concentrations of a two-fold dilution series and incubated on ice for at least 30 minutes. PBS-Tween 20 (0.05%) pH 7.5 was used as a dilution buffer. Before the measurement, 3.5 µL of PBS-Tween 20 (0.05%) pH 7.5 was transferred to each of the 24 flow buffer ports of the Fluidity-One M chip plate, and the microfluidic circuits were allowed to prime for at least 1 minute. Then, 2 times 3.5 µL of the 12 different RBD–antibody mixtures were transferred to the 24 sample ports of the Fluidity One-M chip plate to measure a binding curve of 12 antibody concentrations in duplicate. On the Fluidity One-M, the Alexa-647 detection setting and size-range setting of 3 – 14 nm was used. *K*_D_ values were determined by non-linear least squares fitting as described previously^18^ using Prism (GraphPad Software).

### Microfluidic Antibody Affinity Profiling (MAAP) on the working reagent for anti-SARS-CoV-2 immunoglobulin

The working reagent for anti-SARS-CoV-2 immunoglobulin (National Institute for Biological Standards and Control 21/234) is a calibrated product equivalent to the high concentration sample (NIBSC 20/150) from the WHO working standard for anti-SARS-CoV-2 immunoglobulin (NIBSC 20/268). NIBSC 21/234 consist of pooled plasma from individuals who recovered from COVID-19 and was collected between April and May 2020^23^. MAAP was performed as described previously^19–21^. In brief, fluorescently labelled spike RBD from wild type, Delta, and Omicron was mixed with various dilutions of convalescent plasma and incubated on ice for at least 30 min. A buffer containing PBS at pH 7.4, 10% glycerol, and 5% (w/v) human serum albumin was used for plasma dilutions. Equilibrium binding of RBD to plasma antibodies was assessed by monitoring hydrodynamic radii (*R*_h_) on the Fluidity One-M. The measurement protocol was the same as for purified proteins measured in buffer, with the only difference being that 3.5 µL of the plasma sample instead of PBS-Tween (0.05%) was added to the flow buffer ports of the Fluidity One-M chip plate. Bayesian inference was used to determine *K*_D_ and binding site concentrations from the mode of the joint posterior distribution, also as described previously^19–21^.

## Funding Acknowledgements

Institutional core funding by the University of Zurich and the University Hospital of Zurich, as well as Driver Grant 2017DRI17 of the Swiss Personalized Health Network to A.A; funding by grants of Innovation Fund of the University Hospital Zurich (INOV00096), and of the NOMIS Foundation, the Schwyzer Winiker Stiftung, and the Baugarten Stiftung (coordinated by the USZ Foundation, USZF27101) to A.A. and M.E. V.K. was funded by an NIHR fellowship (PD-2016-09-065). T.P.J.K. is grateful for financial support by the Biotechnology and Biological Sciences Research Council and the European Research Council.

## Author Contributions

S. Fiedler: conceptualization, data curation, formal analysis, supervision, investigation, visualization, methodology, project administration and writing – original draft, review and editing.

S.R.A. Devenish: conceptualization, data curation, formal analysis, supervision, investigation, visualization, methodology, project administration and writing – original draft, review and editing.

A.S. Morgunov: conceptualization, data curation, formal analysis, supervision, investigation, visualization, methodology, project administration and writing – original draft, review and editing.

A. Ilsley: data curation and writing – review and editing.

F. Ricci: data curation and writing – review and editing.

M. Emmenegger: data curation and writing –review and editing.

V. Kosmoliaptsis: conceptualization, writing – review.

A. Lagerwaard: data curation, project administration and writing – review.

J.R. Mills: conceptualization, resources, data curation, funding acquisition and writing – review.

A.M. Sholukh: conceptualization, data curation, investigation, visualization, project administration and writing – original draft, review and editing.

A. Aguzzi: conceptualization, resources, supervision, and writing—original draft, review, and editing.

A. Iwasaki: conceptualization, supervision, and writing—original draft, review, and editing.

A.K. Lynn: conceptualization, resources, data curation, visualization, project administration, funding acquisition and writing – review and editing.

T.P.J. Knowles: conceptualization, resources, supervision, funding acquisition, methodology, and writing—original draft, review, and editing.

## Supplementary Information

**Table S1.**
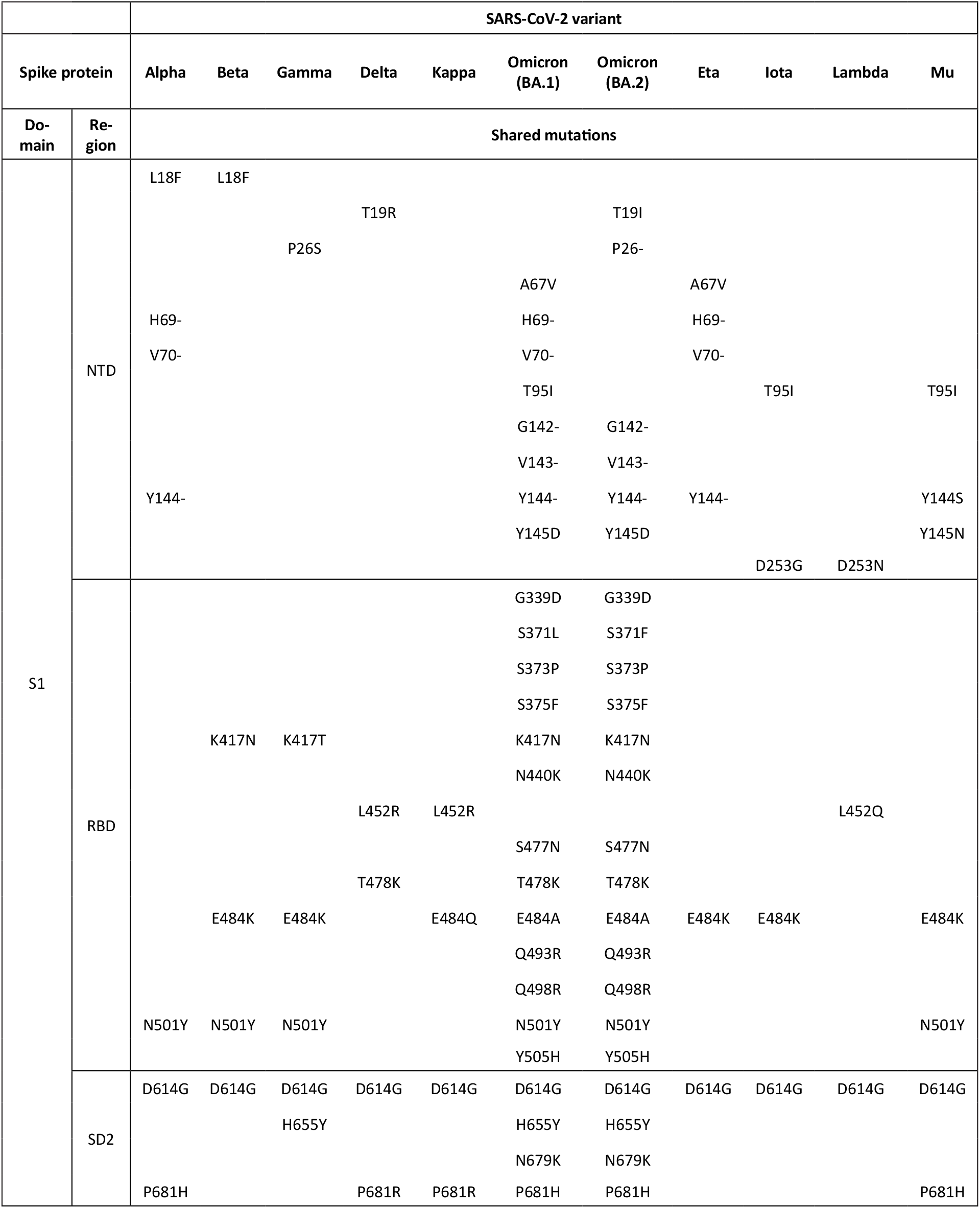
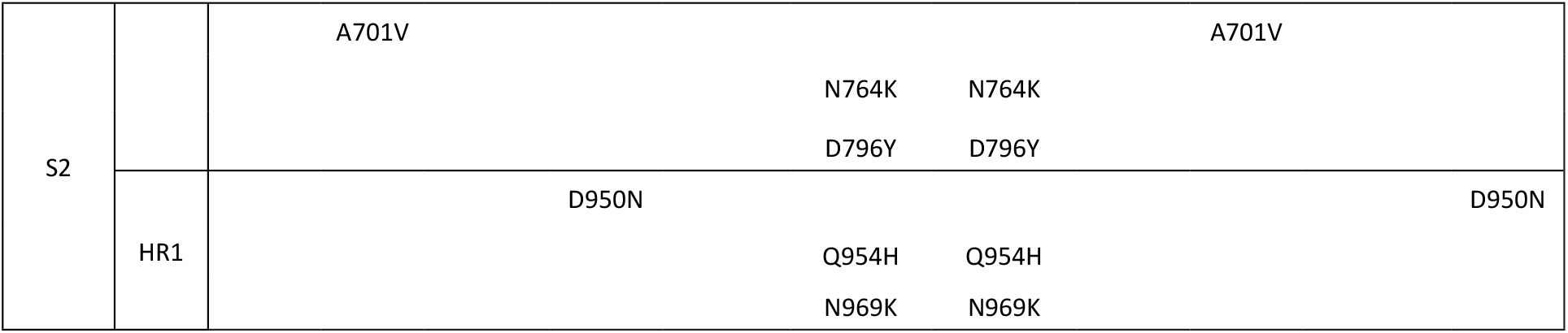
Mutations within the spike protein shared among SARS-CoV-2 variants^32^.

**Table S2.**
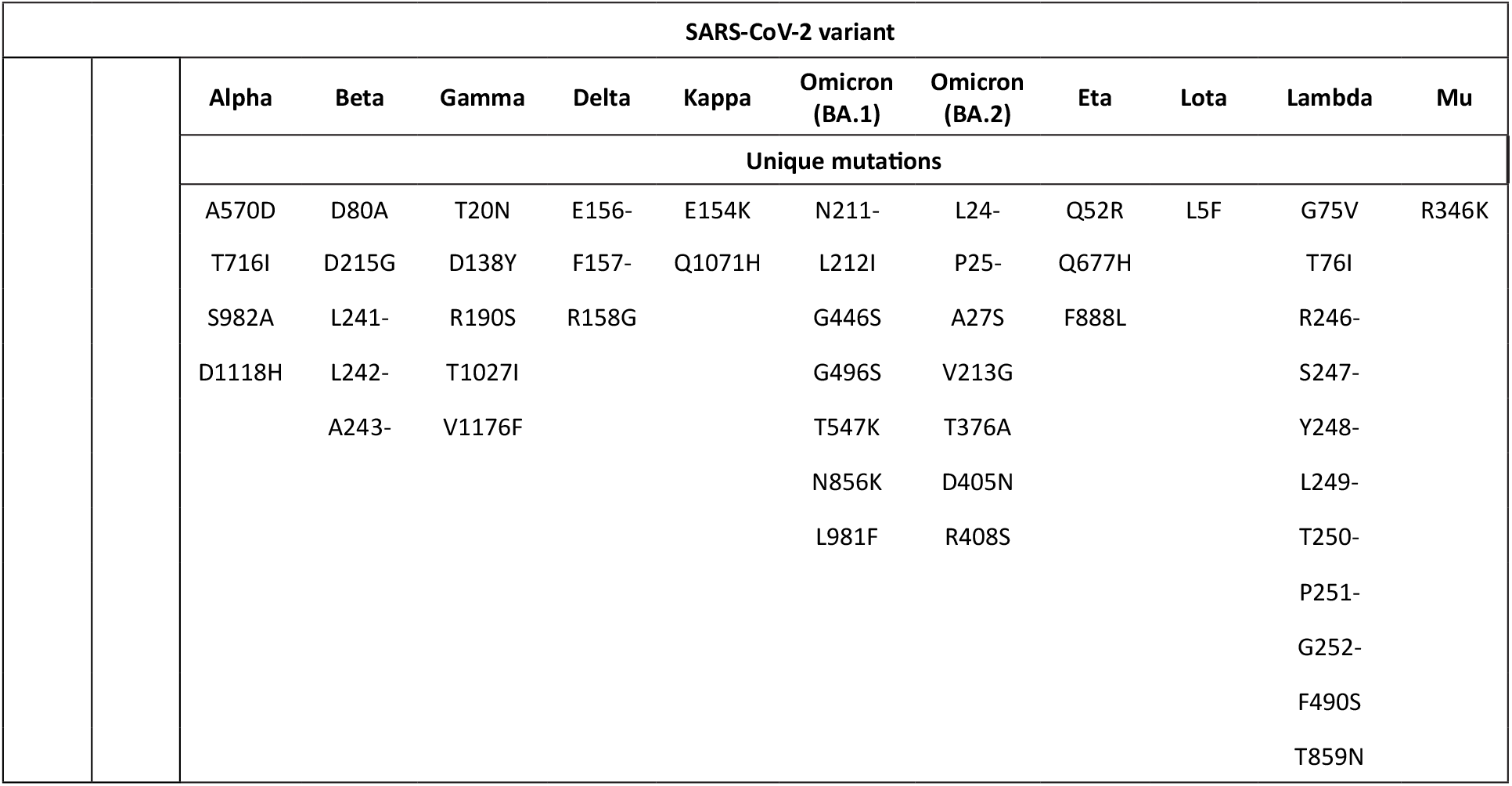
Mutations within the spike protein unique among SARS-CoV-2 variants^32^.

**Table S3.**
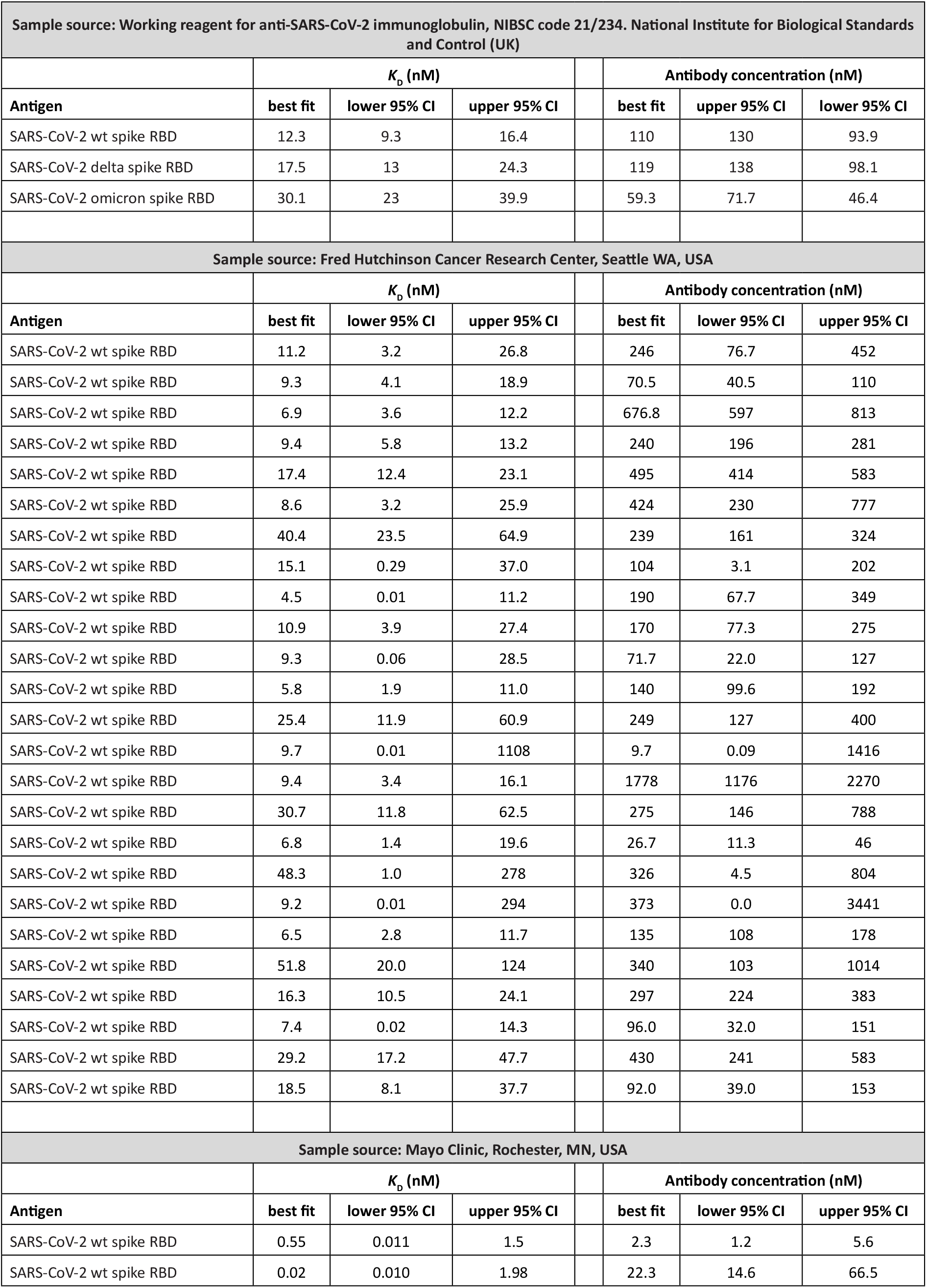
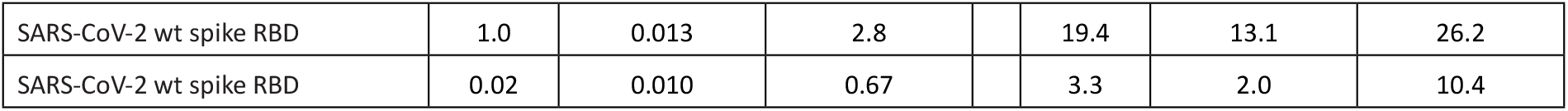
Summary of unpublished microfluidic antibody affinity profiling data from SARS-CoV-2 convalescent samples shown in Figure 3. Results from samples collected by Blutspendedienst (BDS) Kanton Zürich and University Hospital Zurich (CH) have been published previously in Life Sci. Alliance 2021, 5 (2), e202101270. Results from samples purchased from BioIVT LLC, have been published previously on bioRxiv (doi: 10.1101/2021.07.23.453352 and doi: 10.1101/2021.07.23.453327).

