## Supplementary Information for "Serological fingerprints link antiviral activity of therapeutic antibodies to affinity and concentration" (from another version of this preprint: version 1, 2022-02-04)

### Antibody affinity and concentration of convalescent sera provide context for reduced SARS-CoV-2 Omicron spike affinity of therapeutic antibodies

Sebastian Fiedler[1], Sean R. A. Devenish[1], Alexey S. Morgunov[1,2], Alison Ilsley[1], Francesco Ricci[1], Marc Emmenegger[3], Vasilis Kosmoliaptsis[4,5,6], Elitza S. Theel[7], John R. Mills[8,9], Anton M. Sholukh[10], Adriano Aguzzi[3], Akiko Iwasaki[11,12,13,14], Andrew K. Lynn[1], Tuomas P. J. Knowles[1,2,15*]

[1]Fluidic Analytics, Unit A, The Paddocks Business Centre, Cherry Hinton Road, Cambridge CB1 8DH, United Kingdom

[2]Centre for Misfolding Diseases, Yusuf Hamied Department of Chemistry, University of Cambridge, Lensfield Road, Cambridge CB2 1EW, United Kingdom

[3]Institute of Neuropathology, University of Zurich, 8091 Zurich, Switzerland

[4]Department of Surgery, University of Cambridge, Addenbrookes Hospital, Cambridge, CB2 0QQ, United Kingdom

[5]NIHR Blood and Transplant Research Unit in Organ Donation and Transplantation, University of Cambridge, Hills Road, Cambridge CB2 0QQ, United Kingdom

[6]NIHR Cambridge Biomedical Research Centre, Hills Road, Cambridge CB2 0QQ, United Kingdom

[7]Division of Clinical Microbiology, Department of Laboratory Medicine and Pathology, Mayo Clinic, Rochester, Minnesota, USA

[8]Department of Laboratory Medicine and Pathology, Mayo Clinic, Rochester, Minnesota, USA

[9]Center for MS and Autoimmune Neurology, Mayo Clinic, Rochester, Minnesota, USA

[10]Vaccine and Infectious Disease Division, Fred Hutchinson Cancer Research Center, Seattle, WA 98109, USA

[11]Department of Immunobiology, Yale School of Medicine, New Haven, CT 06519, USA

[12]Department of Epidemiology of Microbial Diseases, Yale School of Public Health, New Haven, CT 06510, USA

[13]Department of Molecular, Cellular and Developmental Biology, Yale University, New Haven, CT 06511, USA

[14]Howard Hughes Medical Institute, Chevy Chase, MD 20815, USA

[15]Cavendish Laboratory, Department of Physics, University of Cambridge, JJ Thomson Ave, Cambridge CB3 0HE, United Kingdom

**Table S1.** Mutations within the spike protein shared among SARS-CoV-2 variants[32].

| | | SARS-CoV-2 variant | | | | | | | | | | |
|---|---|---|---|---|---|---|---|---|---|---|---|---|
| **Spike protein** | | **Alpha** | **Beta** | **Gamma** | **Delta** | **Kappa** | **Omicron (BA.1)** | **Omicron (BA.2)** | **Eta** | **Iota** | **Lambda** | **Mu** |
| **Do-main** | **Re-gion** | **Shared mutations** | | | | | | | | | | |
| S1 | NTD | L18F | L18F | | | | | | | | | |
| | | | | | T19R | | | T19I | | | | |
| | | | | P26S | | | | P26- | | | | |
| | | | | | | | A67V | | A67V | | | |
| | | H69- | | | | | H69- | | H69- | | | |
| | | V70- | | | | | V70- | | V70- | | | |
| | | | | | | | T95I | | | T95I | | T95I |
| | | | | | | | G142- | G142- | | | | |
| | | | | | | | V143- | V143- | | | | |
| | | Y144- | | | | | Y144- | Y144- | Y144- | | | Y144S |
| | | | | | | | Y145D | Y145D | | | | Y145N |
| | | | | | | | | | | D253G | D253N | |
| | RBD | | | | | | G339D | G339D | | | | |
| | | | | | | | S371L | S371F | | | | |
| | | | | | | | S373P | S373P | | | | |
| | | | | | | | S375F | S375F | | | | |
| | | | K417N | K417T | | | K417N | K417N | | | | |
| | | | | | | | N440K | N440K | | | | |
| | | | | | L452R | L452R | | | | | L452Q | |
| | | | | | | | S477N | S477N | | | | |
| | | | | | T478K | | T478K | T478K | | | | |
| | | | E484K | E484K | | E484Q | E484A | E484A | E484K | E484K | | E484K |
| | | | | | | | Q493R | Q493R | | | | |
| | | | | | | | Q498R | Q498R | | | | |
| | | N501Y | N501Y | N501Y | | | N501Y | N501Y | | | | N501Y |
| | | | | | | | Y505H | Y505H | | | | |
| | SD2 | D614G | D614G | D614G | D614G | D614G | D614G | D614G | D614G | D614G | D614G | D614G |
| | | | | H655Y | | | H655Y | H655Y | | | | |
| | | | | | | | N679K | N679K | | | | |
| | | P681H | | | P681R | P681R | P681H | P681H | | | | P681H |
| S2 | | | A701V | | | | | | | A701V | | |
| | | | | | | | N764K | N764K | | | | |
| | | | | | | | D796Y | D796Y | | | | |
| | HR1 | | | | D950N | | | | | | | D950N |
| | | | | | | | Q954H | Q954H | | | | |
| | | | | | | | N969K | N969K | | | | |

**Table S2.** Mutations within the spike protein unique among SARS-CoV-2 variants[32].

| SARS-CoV-2 variant | | | | | | | | | | | | |
|---|---|---|---|---|---|---|---|---|---|---|---|---|
| | | **Alpha** | **Beta** | **Gamma** | **Delta** | **Kappa** | **Omicron (BA.1)** | **Omicron (BA.2)** | **Eta** | **Lota** | **Lambda** | **Mu** |
| | | **Unique mutations** | | | | | | | | | | |
| | | A570D | D80A | T20N | E156- | E154K | N211- | L24- | Q52R | L5F | G75V | R346K |
| | | T716I | D215G | D138Y | F157- | Q1071H | L212I | P25- | Q677H | | T76I | |
| | | S982A | L241- | R190S | R158G | | G446S | A27S | F888L | | R246- | |
| | | D1118H | L242- | T1027I | | | G496S | V213G | | | S247- | |
| | | | A243- | V1176F | | | T547K | T376A | | | Y248- | |
| | | | | | | | N856K | D405N | | | L249- | |
| | | | | | | | L981F | R408S | | | T250- | |
| | | | | | | | | | | | P251- | |
| | | | | | | | | | | | G252- | |
| | | | | | | | | | | | F490S | |
| | | | | | | | | | | | T859N | |

**Table S3**. Summary of unpublished microfluidic antibody affinity profiling data from SARS-CoV-2 convalescent samples shown in Figure 3. Results from samples collected by Blutspendedienst (BDS) Kanton Zürich and University Hospital Zurich (CH) have been published previously in Life Sci. Alliance 2021, 5 (2), e202101270. Results from samples purchased from BioIVT LLC, have been published previously on bioRxiv (doi: 10.1101/2021.07.23.453352 and doi: 10.1101/2021.07.23.453327).

| **Sample source: Working reagent for anti-SARS-CoV-2 immunoglobulin, NIBSC code 21/234. National Institute for Biological Standards and Control (UK)** | | | | | | | |
|---|---|---|---|---|---|---|---|
| | **$K_D$ (nM)** | | | | **Antibody concentration (nM)** | | |
| **Antigen** | **best fit** | **lower 95% CI** | **upper 95% CI** | | **best fit** | **upper 95% CI** | **lower 95% CI** |
| SARS-CoV-2 wt spike RBD | 12.3 | 9.3 | 16.4 | | 110 | 130 | 93.9 |
| SARS-CoV-2 delta spike RBD | 17.5 | 13 | 24.3 | | 119 | 138 | 98.1 |
| SARS-CoV-2 omicron spike RBD | 30.1 | 23 | 39.9 | | 59.3 | 71.7 | 46.4 |
| **Sample source: Fred Hutchinson Cancer Research Center, Seattle WA, USA** | | | | | | | |
| | **$K_D$ (nM)** | | | | **Antibody concentration (nM)** | | |
| **Antigen** | **best fit** | **lower 95% CI** | **upper 95% CI** | | **best fit** | **lower 95% CI** | **upper 95% CI** |
| SARS-CoV-2 wt spike RBD | 11.2 | 3.2 | 26.8 | | 246 | 76.7 | 452 |
| SARS-CoV-2 wt spike RBD | 9.3 | 4.1 | 18.9 | | 70.5 | 40.5 | 110 |
| SARS-CoV-2 wt spike RBD | 6.9 | 3.6 | 12.2 | | 676.8 | 597 | 813 |
| SARS-CoV-2 wt spike RBD | 9.4 | 5.8 | 13.2 | | 240 | 196 | 281 |
| SARS-CoV-2 wt spike RBD | 17.4 | 12.4 | 23.1 | | 495 | 414 | 583 |
| SARS-CoV-2 wt spike RBD | 8.6 | 3.2 | 25.9 | | 424 | 230 | 777 |
| SARS-CoV-2 wt spike RBD | 40.4 | 23.5 | 64.9 | | 239 | 161 | 324 |
| SARS-CoV-2 wt spike RBD | 15.1 | 0.29 | 37.0 | | 104 | 3.1 | 202 |
| SARS-CoV-2 wt spike RBD | 4.5 | 0.01 | 11.2 | | 190 | 67.7 | 349 |
| SARS-CoV-2 wt spike RBD | 10.9 | 3.9 | 27.4 | | 170 | 77.3 | 275 |
| SARS-CoV-2 wt spike RBD | 9.3 | 0.06 | 28.5 | | 71.7 | 22.0 | 127 |
| SARS-CoV-2 wt spike RBD | 5.8 | 1.9 | 11.0 | | 140 | 99.6 | 192 |
| SARS-CoV-2 wt spike RBD | 25.4 | 11.9 | 60.9 | | 249 | 127 | 400 |
| SARS-CoV-2 wt spike RBD | 9.7 | 0.01 | 1108 | | 9.7 | 0.09 | 1416 |
| SARS-CoV-2 wt spike RBD | 9.4 | 3.4 | 16.1 | | 1778 | 1176 | 2270 |
| SARS-CoV-2 wt spike RBD | 30.7 | 11.8 | 62.5 | | 275 | 146 | 788 |
| SARS-CoV-2 wt spike RBD | 6.8 | 1.4 | 19.6 | | 26.7 | 11.3 | 46 |
| SARS-CoV-2 wt spike RBD | 48.3 | 1.0 | 278 | | 326 | 4.5 | 804 |
| SARS-CoV-2 wt spike RBD | 9.2 | 0.01 | 294 | | 373 | 0.0 | 3441 |
| SARS-CoV-2 wt spike RBD | 6.5 | 2.8 | 11.7 | | 135 | 108 | 178 |
| SARS-CoV-2 wt spike RBD | 51.8 | 20.0 | 124 | | 340 | 103 | 1014 |
| SARS-CoV-2 wt spike RBD | 16.3 | 10.5 | 24.1 | | 297 | 224 | 383 |
| SARS-CoV-2 wt spike RBD | 7.4 | 0.02 | 14.3 | | 96.0 | 32.0 | 151 |
| SARS-CoV-2 wt spike RBD | 29.2 | 17.2 | 47.7 | | 430 | 241 | 583 |
| SARS-CoV-2 wt spike RBD | 18.5 | 8.1 | 37.7 | | 92.0 | 39.0 | 153 |
| **Sample source: Mayo Clinic, Rochester, MN, USA** | | | | | | | |
| | **$K_D$ (nM)** | | | | **Antibody concentration (nM)** | | |
| **Antigen** | **best fit** | **lower 95% CI** | **upper 95% CI** | | **best fit** | **lower 95% CI** | **upper 95% CI** |
| SARS-CoV-2 wt spike RBD | 0.55 | 0.011 | 1.5 | | 2.3 | 1.2 | 5.6 |
| SARS-CoV-2 wt spike RBD | 0.02 | 0.010 | 1.98 | | 22.3 | 14.6 | 66.5 |

| | | | | | | | |
|---|---|---|---|---|---|---|---|
| SARS-CoV-2 wt spike RBD | 1.0 | 0.013 | 2.8 | | 19.4 | 13.1 | 26.2 |
| SARS-CoV-2 wt spike RBD | 0.02 | 0.010 | 0.67 | | 3.3 | 2.0 | 10.4 |
